# Early life stress affects the miRNA cargo in epididymal extracellular vesicles in mouse

**DOI:** 10.1101/2021.04.29.441964

**Authors:** Anar Alshanbayeva, Deepak K. Tanwar, Martin Roszkowski, Francesca Manuella, Isabelle M. Mansuy

## Abstract

Sperm RNA can be modified by environmental factors and has been implicated in communicating signals about changes in a father’s environment to the offspring. The RNA composition of sperm is influenced during its final stage of maturation in the epididymis by extracellular vesicles released by epididymal cells. We studied the effect of exposure to stress in postnatal life on the transcriptome of epididymal extracellular vesicles using a mouse model of transgenerational transmission. We found that the small RNA signature of epididymal extracellular vesicles, particularly miRNAs, is altered in adult males exposed to postnatal stress. miRNAs changes correlate with differences in the expression of their target genes in sperm and zygotes generated from that sperm. These results suggest that stressful experiences in early life can have persistent biological effects on the male reproductive tract that may in part be responsible for the transmission of the effects of exposure to the offspring.

**Summary Sentence:** miRNA cargo of extracellular vesicles in cauda epididymis is changed by paternal exposure to early life stress, which correlates with differences in the expression of their target genes in sperm and zygotes generated from that sperm

## Introduction

Post-testicular maturation of spermatozoa in the epididymis is an elaborate process that involves modifications of sperm RNA, protein and lipid content [1–5]. The epididymis is segmented into different parts, including the initial segment, caput, corpus and cauda. Each segment has a distinct gene expression profile, and different protein and lipid composition. Some modifications in epididymal spermatozoa are conserved across species [6]. For example, approximately 50% of miRNAs, a class of small RNAs that are modified during caput to cauda epididymis transit, is identical in mouse and bovine spermatozoa [6]. One mechanism by which small RNA load in spermatozoa is modified along the epididymis is by uptake of extracellular vesicles (EVs), also known as epididymosomes, which are produced by epididymal epithelial cells [7]. Studies have shown that epididymosomes can be taken up by maturing sperm through proteins present on the sperm head such as dynamin in mice and tetraspanins or syntenins in humans [7–10]. Co-incubation experiments provided evidence for epididymosome-mediated transfer of miRNAs to spermatozoa [7]. Exogenous DNA and RNA can also be directly taken up by spermatozoa via artificial liposomes [11].

However, it is still not clear if changes in small RNA composition of spermatozoa occurring during epididymal transit are required for embryonic development, and studies on the subject have been conflicting [12,13]. Changes in sperm small RNA have nevertheless been suggested to play a role in the transmission of information about paternal experiences to the progeny and can influence their developmental trajectory [2,14,15]. Epididymosomal small RNA content can also be altered by exposure, for instance, to dietary insult or stress [2,14]. For instance, epididymosomal miRNAs are changed by exposure to chronic stress [14] and low-protein diet [2] in mice.

Transmission of information about paternal exposure to the offspring depends on the type of exposure, as well as on its duration and developmental window [16]. To date, little is known about the long-term effects of early life stress, particularly stress experienced after birth on epididymosomal small RNA composition in adulthood and whether any change can influence gene expression in sperm and in zygotes generated from that sperm. Using a transgenerational mouse model of postnatal stress (based on unpredictable maternal separation combined with unpredictable maternal stress, MSUS) [17], we show that the miRNA signature of cauda epididymosomes in adult males is altered by exposure to postnatal stress. The alterations in miRNA composition are found to correlate with changes in the expression of targets of these miRNAs in sperm and zygotes.

## Results

### Isolation of cauda epididymosomes confirmed by several methods

To characterize RNA composition of cauda epididymosomes, epididymosomes were first isolated from adult control males and adult males exposed to MSUS by high-speed ultracentrifugation (Figure 1A). Successful isolation of cauda epididymosomes was confirmed by electron microscopy, immunoblotting and nanoparticle-tracking analyses (Figure 1). The presence and purity of epididymosomes was further validated by staining with the EV-specific marker CD9 and absence of the cellular marker GAPDH (Figure 1B). Size analysis by nanoparticle-tracking indicated that the collected particles are 50-300 nm in diameter (Figure 1D), and imaging by transmission electron microscopy showed the typical cup-shaped structures of epididymosomes (Figure 1C, Supplementary Figure 1A) [18]. RNA profiling by high-resolution automated electrophoresis showed enrichment for small RNAs of different lengths, similar to previous studies on cauda epididymosomal RNA content (Supplementary Figure 1B) [2,13].

**Figure 1.**
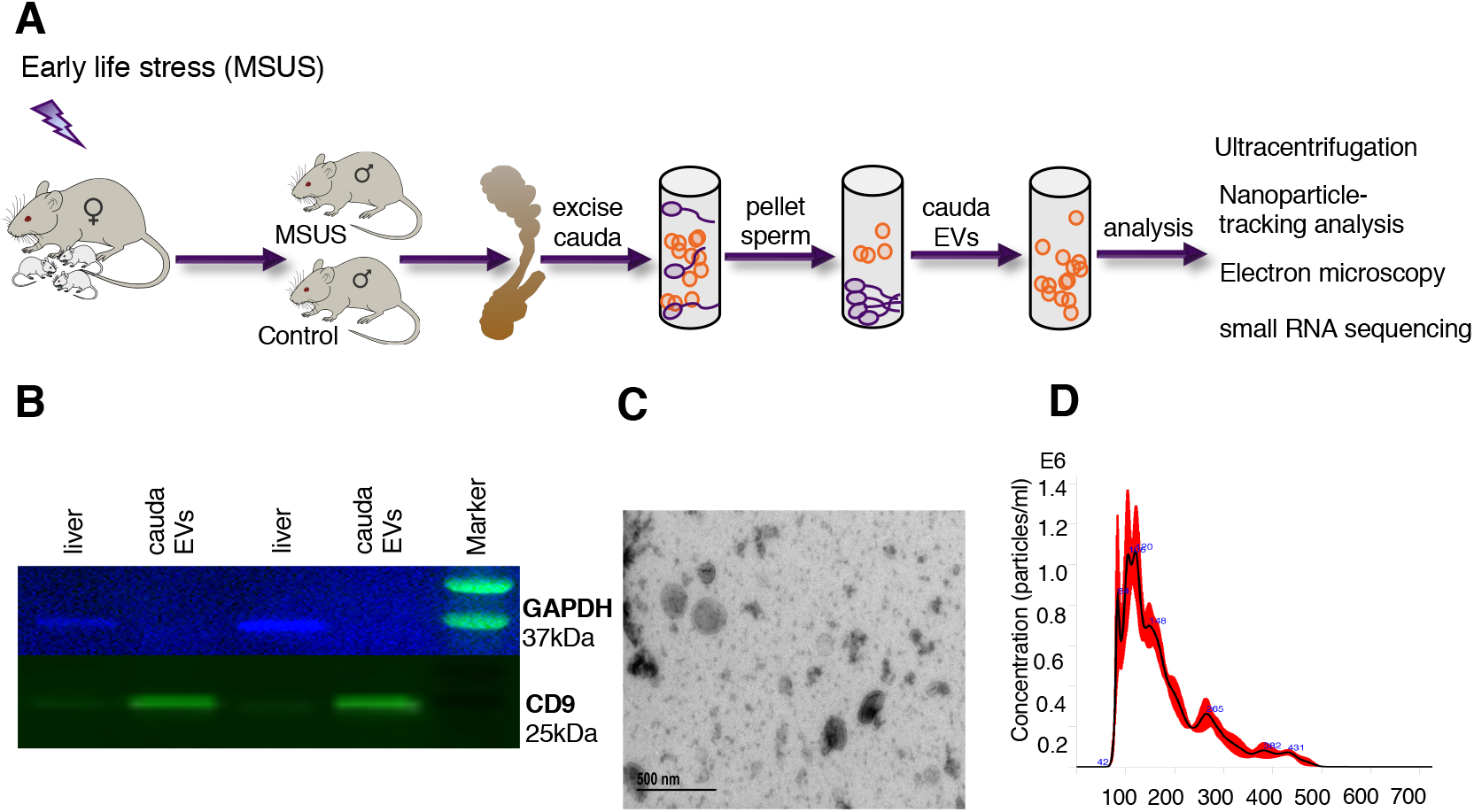
Isolation and characterization of cauda epididymosomes. **(A)** Schematic representation of cauda epididymosomes preparation. **(B)** Immunoblot analysis was used to confirm the purity of isolations by staining with epididymosomal marker CD9 and absence of cellular marker GAPDH in the ultracentrifuged pellet. (**C**) Electron microscopy images of the preparations were used to access the size and heterogeneity of the isolated populations. (D) Nanoparticle-tracking analysis by dynamic light scattering showed isolation of particles of expected size of 50-300 nm.

### The number and size of epididymosomes in adult males are not altered by postnatal stress

We next examined the number and size of epididymosomes in adult MSUS and control males by dynamic light scattering. No significant difference in the number or mean size of cauda epididymosomes could be detected between MSUS and control males (Figure 2A, 2B). Since most epididymosomal secretion occurs via apocrine secretion from principal cells located in caput epididymis, we also examined the level of expression of genes involved in extracellular vesicles secretion. We chose Ras-related protein Rab-5A (*Rab5*) and Ras-related protein Rab-7A (*Rab7*), which are involved in vesicle trafficking, the SNARE family protein vesicle-associated membrane protein 7 (*Vamp7*) and SNARE recognition molecule synaptobrevin homolog YKT6 (*Ykt6*), involved in vesicle fusion. No significant change in the expression of these genes could be detected in caput epididymis between MSUS and control mice, despite a consistent trend for decreased expression in MSUS mice (Figure 2C).

**Figure 2.**
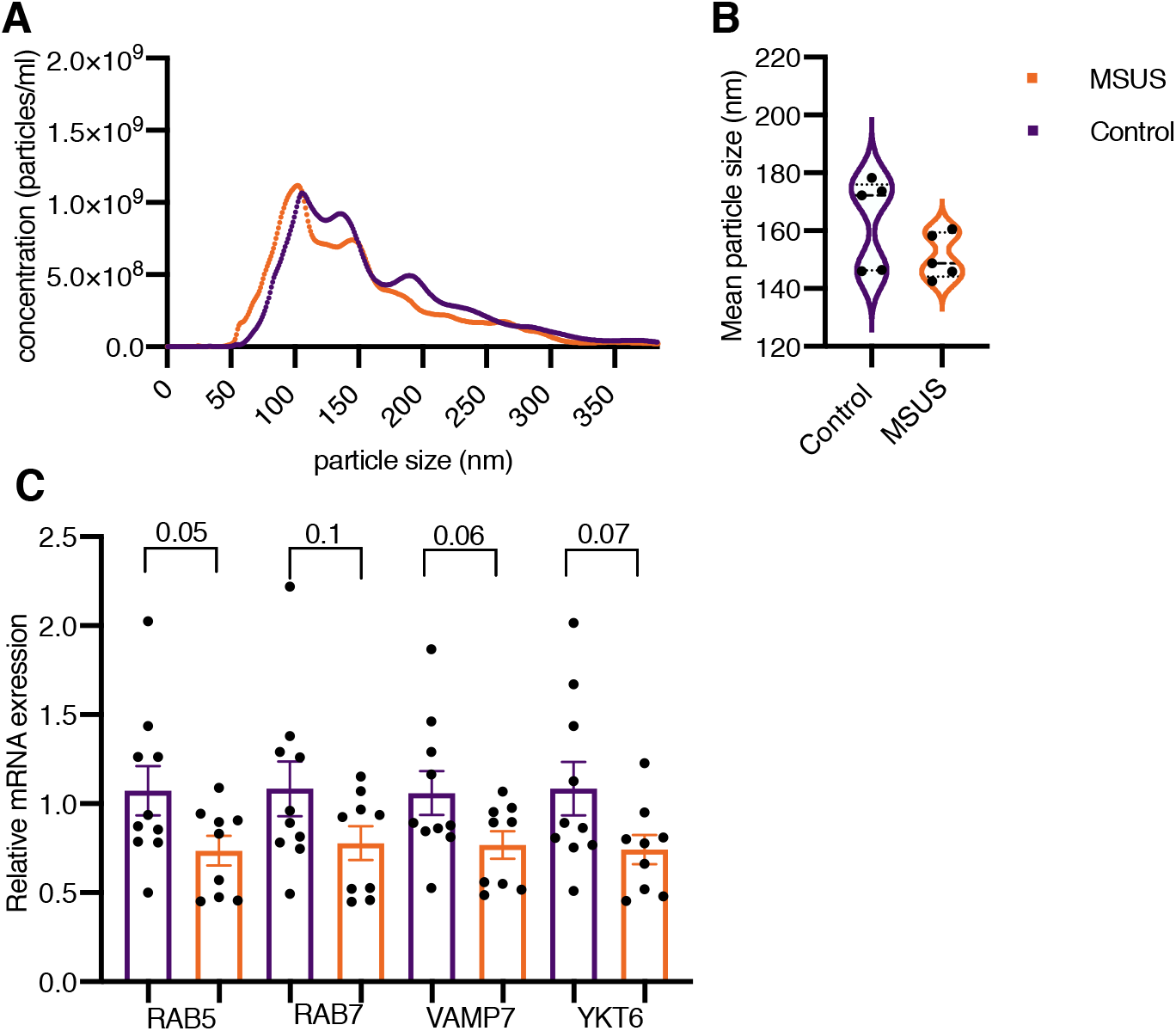
Comparison of epididymosomal number, size and release machinery in MSUS and control. **(A)** Nanoparticle-tracking analysis showed no difference in the number of cauda epididymosomes between MSUS and control males. The plots were generated from average values across replicates (N = 5 animals/group). Data are presented as mean ± standard error of the mean (SEM). *P* < 0.05. (**B**) Quantification of nanoparticle-tracking analysis showed the mean size of cauda epididymosomes was not changed between the two groups (N = 5 animals/group). Data are presented as mean ± SEM. *P* < 0.05. (**C**) Expression of genes involved in vesicular secretion in the caput epididymis from adult males measured by qRT-PCR. The experiments were performed in triplicates without pooling (N = 8 animals/group). Expression of Gapdh was used as endogenous control to normalize the expression level of the target genes. Data are presented as mean ± SEM. *P* < 0.05.

### miRNAs are persistently altered by postnatal stress in cauda epididymosomes

Epididymosomal small RNAs are known to be affected by changing environmental conditions in rodents. Small RNAs, like tRNA-derived fragments (tRFs) are believed to act as messengers of a father’s experiences to the offspring [2,14]. Our previous work showed that early postnatal stress alters small and long RNA content in sperm in adult males [15]. Since caudal sperm and epididymosomal small RNA profiles are highly similar [2], we examined whether small RNA content of cauda epididymosomes is also altered by postnatal stress. RNA of different sizes was observed in cauda epididymosomes, with the majority of small RNA reads mapping to tRNAs as previously observed (Figure 3A) [2]. When plotting the top different small RNAs between the two groups, the majority of small RNA differences between MSUS and control epididymosomes appeared to be in miRNAs (Figure 3B). Therefore, we performed size-selection on the same libraries and re-sequenced them to enrich for miRNAs (Supplementary Figure 2A, 2C). As expected, size-selection did not alter the abundance of different miRNAs and uniformly enriched the miRNA fraction in all samples (Supplementary Figure 2B, 2C, 2D). Pathway analysis of small RNA-sequencing (sRNA-seq) datasets after size-selection revealed that the most up-regulated and down-regulated miRNAs (*P* < 0.05) in MSUS cauda epididymosomes have target mRNAs that encode proteins involved in pathways, including fatty acid metabolism, steroid biosynthesis, lysine degradation and thyroid hormone signaling (Figure 3C). Notably, similar pathways are altered in plasma of MSUS males as shown by unbiased metabolomic analysis that we previously conducted [19]. In particular, metabolites implicated in polyunsaturated fatty acid biogenesis were up-regulated, whereas steroidogenesis and the steroidogenic ligand aldosterone were down-regulated in circulation of MSUS males [19]. These results suggest systemic changes in fatty acid metabolism and steroidogenesis pathways in adult MSUS males. Notably, steroidogenesis was already altered during postnatal life in MSUS males with total cholesterol significantly decreased in testis, while HDL cholesterol was significantly increased in liver in MSUS males at postnatal day 28 (Figure 3D, 3F). Since the primary role of HDL cholesterol in blood is to transport excess cholesterol from peripheral tissues and lipoproteins to the liver, an increase in HDL in the liver corresponds to the HDL decrease in testis. However, cholesterol was no longer altered in testis of adult MSUS males (Figure 3E), suggesting a transient alteration. The androgen receptor, that binds androgens which are derived from cholesterol, showed however a trend for decreased expression in adult caput epididymis (Figure 3G), suggesting potential secondary effects of lower cholesterol in testis earlier in life.

**Figure 3.**
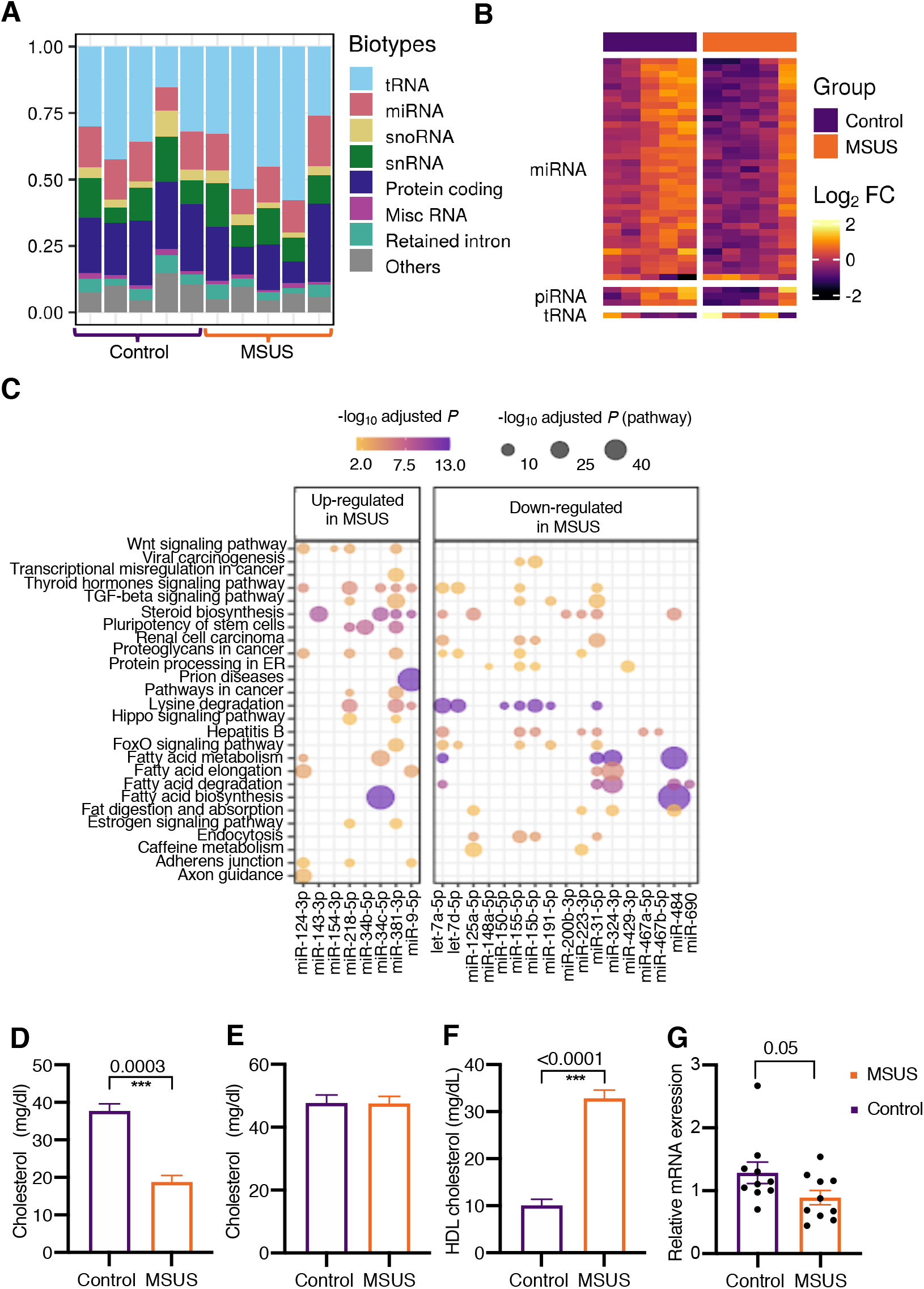
Target pathways of up- and down-regulated miRNAs in MSUS cauda epididymosomes. Alterations in steroidogenesis in MSUS males. **(A)** Representative distribution of RNA biotypes from cauda epididymosomal sequencing (N = 10 animals, 5 animals/group). (**B**) Heatmap of the most abundant small RNAs (n = 39). Expression fold-change (log2 FC) was calculated by subtracting log2 counts per million (CPM) of MSUS from Controls. Each row depicts a small RNA and each column depicts a sample. Samples and RNAs are ordered by “PCA” method using seriation (R package). (**C**) Dot plot of miRNAs and pathways. Color-scale of the dot represents -log_10_ adjusted *P* of miRNA in a pathway and size of the dot represents -log_10_ adjusted *P* of the pathway. Total cholesterol measurements in whole testes of males at postnatal day 28 (N = 4 males/group) (**D**) or adulthood (N = 10 Controls, N = 7 MSUS) (**E**). (**F**) HDL cholesterol level in the liver of males at postnatal day 28 (N = 6 males/group). (**G**) Relative expression level of the androgen receptor in adult caput epididymis measured by qRT-PCR. qRT-PCR experiments were performed in triplicates, without pooling (N = 10 animals/group). (**D, E, F, G**) Data are presented as mean ± standard error of the mean (SEM). *P* < 0.05.

### mRNA targets of miRNAs from cauda epididymosomes are altered by postnatal stress in sperm and zygotes

The relative abundance of miRNAs in cauda epididymosomes and mature sperm was significantly correlated (Figure 4C), consistent with previous results [2]. Since cauda epididymosomes carry small RNA payloads matching those of mature sperm and are part of the ejaculate [20,21], they may contribute to the information delivered to the embryo. Therefore, we looked at the mRNA targets of the miRNAs significantly changed in MSUS cauda epididymosomes (adjusted *P* ≤ 0.05), which include miR-871-5p, miR-31-5p, miR-155-5p and miR34c-5p, among differentially expressed genes we previously identified in MSUS sperm and zygotes (*P* < 0.05) [15,22]. For this, we plotted the cumulative log fold-change distribution of genes versus the different numbers of conserved binding sites for these miRNAs (Figure 4A and 4B, Supplementary Figure 5). Target genes with 3 binding sites for miR-31-5p, a miRNA differentially expressed in MSUS cauda epididymosomes, had increased expression in sperm and decreased expression in zygotes from MSUS males (Figure 4A, 4B). However, not all targets of significantly altered miRNAs from MSUS cauda epididymosomes showed corresponding expression changes in sperm and zygotes (Supplementary Figure 5). We performed miRNA-gene interaction analysis based on experimentally validated data from Tarbase [23]. From this analysis, we identified that the 5 miRNAs significantly changed in MSUS cauda epididymosomes target genes that are part of pathways involved in steroid biosynthesis, ECM-receptor interaction and cell adhesion molecules (Figure 4D).

**Figure 4.**
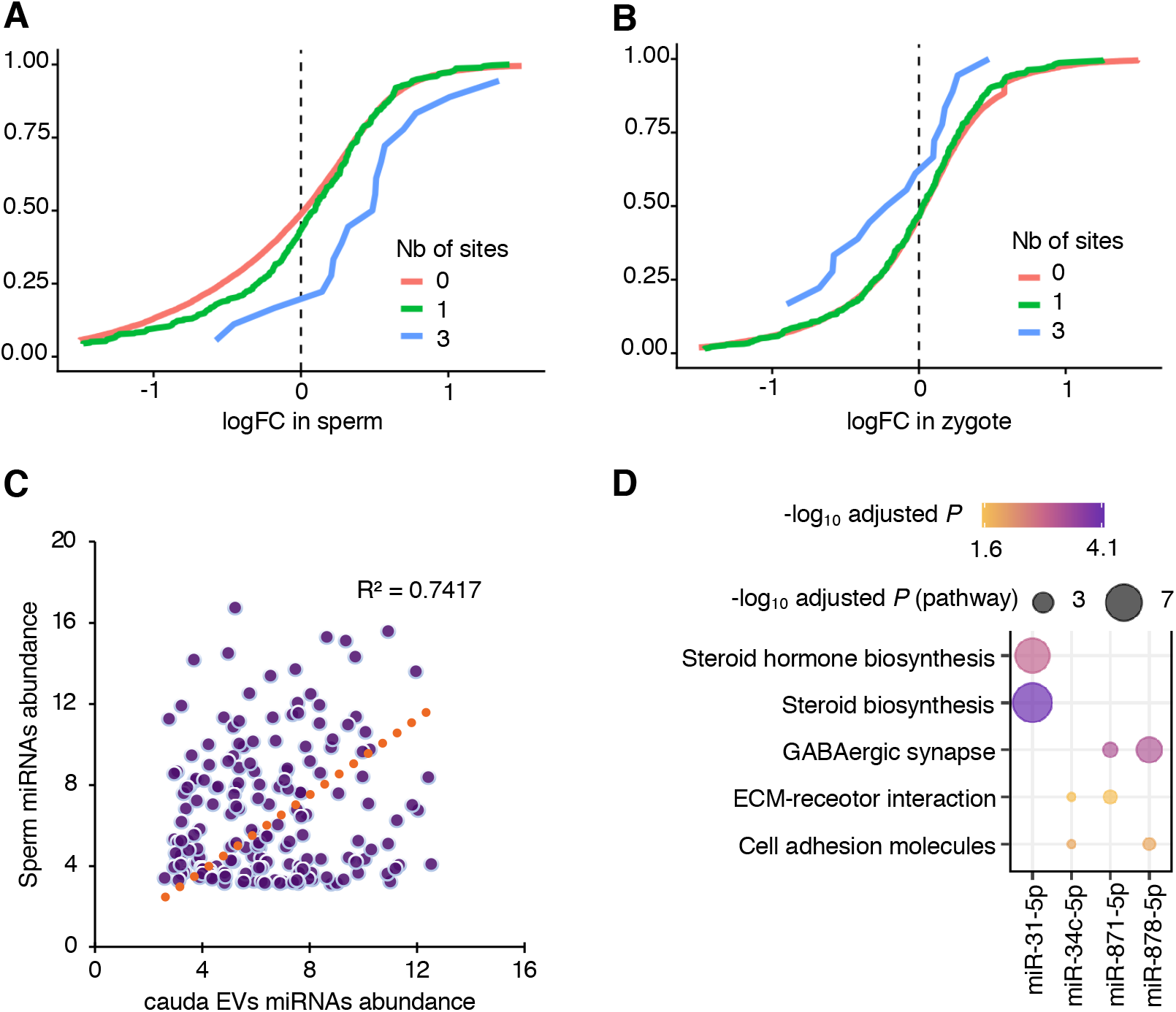
Targets of miRNAs from cauda epididymosomes are altered by postnatal stress in sperm and zygotes. Cumulative distribution plots of miR-31-5p targets in differentially expressed genes (*P* < 0.05) from MSUS sperm RNA-seq (**A**) and zygote RNA-seq (**B**). (**C**) miRNA abundance of sperm plotted against abundance in cauda epididymosomes. Coefficient of determination (R^2^) = 0.74. (**D**) Dot plot of the top target pathways (adjusted *P* < 0.05) of miRNAs differentially expressed (adjusted *P* ≤ 0.05) in MSUS cauda epididymosomes. Color-scale of the dot represents -log_10_ adjusted *P* of miRNA in a pathway and size of the dot represents -log_10_ adjusted *P* of the pathway.

## Discussion

The effects of environmental factors on RNA in the male reproductive tract, in particular, the epididymis have been examined in rodent models. Until now, most models have used invasive exposure such as dietary insult and injection of endocrine disruptors and their effects when applied prenatally and sometimes even before conception. Very few studies have examined the effects of non-invasive psychological/emotional exposure such as stress in early life and the effects on epididymal RNA in adulthood. This study addresses the question of whether postnatal stress affects RNA in the epididymis and whether this has consequences for sperm in adulthood and zygotes generated from that sperm.

Using a transgenerational mouse model of early postnatal stress, we show that several miRNAs including miR-871-3p, miR-31-5p, miR-155-5p, miR-878-5p and miR-34c-5p are altered in the epididymis of adult males exposed to stress and that the targets of some of these miRNAs are affected in mature sperm and zygotes. Particularly, miR-31-5p is significantly decreased in cauda epididymosomes from exposed males and its target genes are increased in sperm. Interestingly, these genes are decreased in zygotes generated from that sperm (Figure 4B), suggesting a possible over-compensation during early development. This may also be due to the heterogeneity of epididymosomes which have different size, biogenesis and cellular targeting [24], leading to a dissociation between the RNA content of epididymosomes and transcriptional changes in the zygotes (Figure 4A, 4B and Supplementary Figure 5). It has been suggested that a subset of epididymosomes can communicate with spermatozoa during its epididymal transit [2,7], another subset serves in the communication within epididymal epithelial cells [25], and a third subset is delivered as part of seminal fluid during fertilization [20,21]. Therefore, due to their heterogeneity, not all cauda epididymosomes or their cargo may be delivered to the oocyte upon fertilization, which may explain the differences in miRNAs targets between sperm and zygotes.

Several of the differentially expressed miRNAs in MSUS cauda epididymosomes play a role in metabolic processes and early development [26]. For instance, miR-31-5p is involved in glucose metabolism and fatty acid oxidation [26]. In *A. japonicus*, its target complement C1q Tumor Necrosis Factor-Related Protein 9A (*CTRP9*) protein is negatively associated with the amount of visceral fat and is positively associated with a favourable glucose or metabolic phenotype [27]. Alterations in glucose and insulin metabolism are also characteristic of MSUS mice [15,17]. Other miRNAs significantly changed in MSUS epididymosomes, such as miR-155-5p, which facilitates differentiation of mouse embryonic stem cells, whereas miR-34c-5p is thought to initiate the first embryonic cleavage in mice [26].

The first days after birth are a sensitive period for the development and establishment of tissue-specific niches for several tissues. Epithelial cells, which are the source of epididymosomes in the epididymis, undergo differentiation and expansion postnatally until puberty [28]. Upon completing the expansion, the number of epididymal epithelial cells remains nearly constant in adulthood, thus possibly carrying a memory of early life exposure into adulthood. The postnatal development and differentiation of epididymal epithelial cells primarily depend on testicular signals [28–31]. Since long-term stress affects the coupling of the hypothalamus-pituitary and hypothalamus-gonadal axes, stress-related decrease in steroidogenesis can have profound effects on the differentiation and expansion of epididymal epithelial cells in early postnatal life. A number of studies have shown the importance of the abundance of androgens during postnatal life for epididymal development [29]. Thus, the availability of testicular cholesterol during the differentiation of epididymal epithelial cells has implications for these cells. Systemic alteration in cholesterol metabolism seen in young MSUS males (decreased total cholesterol in testis and increased HDL cholesterol in the liver) may contribute to the metabolic changes seen in adult animals, such as changes in plasma metabolome of steroidogenesis and fatty acid pathways, as well as alterations in glucose and insulin metabolism in adult MSUS males. Moreover, androgen-dependent miRNAs miR-878-5p and miR-871-3p are significantly increased in cauda epididymosomes in MSUS.

In conclusion, our results provide evidence that chronic stress exerted immediately after birth alters miRNAs in extracellular vesicles of the male reproductive tract until adulthood, with effects in mature sperm and zygotes. Such persistent and intergenerational effects *in vivo* point to the sensitivity of the reproductive system to stress exposure and the detrimental consequences for descendants. These consequences likely differ depending on the time window and severity of paternal stress exposure. Further studies will be necessary to determine these effects as well as the source of the vesicles and their cargo miRNAs.

## Materials and Methods

### Animals

Animal experiments were conducted according to the Swiss Law for Animal Protection and were approved by the cantonal veterinary office in Zürich under license number 83/2018. C57Bl/6J mice were obtained from Janvier (France) and bred in-house to generate animals for the experiments. Animals were maintained under a reverse light-dark cycle in a temperature and humidity-controlled environment with constant access to food and water.

### MSUS

To obtain MSUS mice, 3-month old C57Bl/6J dams and their litters were randomly split into MSUS and Control groups. MSUS dams and their litters were subjected to daily 3 hour separation and a stressor as previously described [17], while control dams and pups were left undisturbed. After weaning at postnatal day 21, pups from different dams were randomly assigned to cages of 4-5 mice, in corresponding treatment groups to avoid litter effects.

### Tissue collection

After decapitation and blood collection, mice were pinned on a dissection board and cleaned with alcohol. Epididymis and testis were carefully excised and separated from surrounding adipose tissue. The epididymis was further separated into caput, corpus and cauda. Cauda was excised with several incisions and sperm collected with a swim-up protocol. The supernatant was collected to isolate epididymosomes. The whole testis and caput epididymis were crushed with stainless steel beads in a tissue crusher in cold PBS, centrifuged at 3’000 rcf for 10 min to pellet the tissue and cells and used for total cholesterol and HDL cholesterol measurements.

### Electron microscopy images

Negative staining of cauda epididymosomes was performed with methylcellulose. Briefly, the carrier grid was glow-discharged in plasma for 10 min and washed with a drop of PBS, then incubated in 1% glutaraldehyde (GA) in water for 5 min and washed with water 5 times for 2 min each. Afterwards the grid was incubated in 1% UAc (uranyl acetate) for 5 min and then kept on ice in Methylcellulose/UAc (900 ul Methylcellulose 2 % and 100 ul 3 % UAc) solution. After incubation with Methylcellulose/UAc, the excess liquid was removed by dipping onto a filter paper. The grid was air-dried on ice for 5 min. Imaging was performed with a Transmission Electron Microscope.

### Epididymosome isolation by ultracentrifugation and density-gradient

After pelleting sperm following the sperm swim-up protocol in M2 medium (Sigma, M7167), the supernatant was centrifuged at 2’000 rcf for 10 min, 10’000 rcf for 30 min and then ultracentrifuged at 120’000 rcf at 4 °C for 2 h (TH 64.1 rotor, Thermo Fisher Scientific). The epididymosomal pellet was then washed in PBS at 4 °C and ultracentrifuged at 120’000 rcf at 4 °C for 2 h. The resulting pellet was resuspended in 60 μl of PBS for all downstream analysis.

### Immunoblotting

The PBS-resuspended pellet containing epididymosomes was lysed in 10x RIPA buffer for 5 min at 4 °C. Equal amounts of protein were mixed with 4x Laemmli Sample Buffer (Bio-Rad Laboratories, USA) and loaded on 4-20% Tris-Glycine polyacrylamide gels (Bio-Rad Laboratories, USA). The membranes were blocked in 5% SureBlock (in Tris-buffer with 0.05% Tween-20) for 1 h at room temperature and incubated with primary antibodies overnight at 4 °C with anti-*Cd9* ([1:3000; System Biosciences, Palo Alto, CA, USA] and anti-*Gapdh* [1:5000; Cell Signaling, Davers, MA, USA; 14C10]).

### Nanoparticle tracking analysis

Particle number and size of epididymosomes were measured using a Nanosight NS300 (Malvern, UK) at 20 °C, according to the manufacturer’s instructions and lots were generated using a published method [32]. The following parameters were kept constant for all samples: “Camera level” = 14 and “Detection threshold” = 7. For measurements with Nanosight, the resuspended pellet from ultracentrifugation was diluted to a 1:1000 concentration.

### RNA isolation and epididymosomes profiling

To lyse purified epididymosomes, 33 μl of lysis buffer (6.4 M guanidine HCl, 5 % Tween 20, 5 % Triton, 120 mM EDTA, and 120 mM Tris pH 8.0) per every 60 μl of PBS resuspended pellet was added to each sample, together with 3.3 μl ProteinaseK and 3.3 μl of water. Samples were incubated at 60 °C for 15 min with shaking. 40 μl of water was added and RNA was extracted using the Trizol LS protocol, according to the manufacturer’s instructions. Profiling of the extracted RNA was performed using high-resolution automated electrophoresis on a 2100 Bioanalyzer (Agilent, G2939BA), according to instructions for the RNA 6000 Pico Kit (Agilent, 5067-1513) reagent.

### Preparation and sequencing of small RNA-seq libraries from epididymosomes

sRNA-seq libraries were prepared using the NEB Next Small RNA-sequencing kit (NEB #E7300, New England BioLabs), according to the manufacturer’s instructions. 80-90 ng of total RNA per sample was used to prepare the libraries. The same libraries were sequenced before and after size-selection (target peak 150bp) with the BluePippin System. 200 million reads were obtained for 10 samples, with 125bp single-stranded read-length on a HiSeq2500 sequencer.

### RT-qPCR

For gene expression analysis in caput epididymis, RNA was extracted using the phenol/chloroform extraction method (TRIzol; Thermo Fisher Scientific). Reverse transcription was performed using miScript II RT reagents (Qiagen) -HiFlex buffer, and RT qPCR was performed with QuantiTect SYBR (Qiagen) on the Light-Cycler II 480 (Roche). All samples were run in cycles as follows: 95 °C for 15 min, 45 cycles of 15 sec at 94 °C, 30 sec at 55 °C and 30 sec at 70°C, followed by gradual increase of temperature to 95 °C. The endogenous control *Gapdh* was used for normalization. Primer sequences for individual genes are made available in the Supplementary Table 1. The expression level of genes was analysed using two-tailed Student’s t-test.

### Cholesterol measurements

Testicular and epididymal total cholesterol and HDL cholesterol levels were measured using the CHOL reagent, in conjunction with SYNCHRON LX® System(s), UniCel® DxC 600/800 System(s) and Synchron® Systems Multi Calibrator (Beckman Coulter), according to the manufacturer’s instructions at the Zurich Integrative Rodent Physiology (ZIRP) facility of the University of Zurich.

### Bioinformatics data analysis

Small RNA-sequencing FASTQ files were processed using the ExceRpt pipeline, a previously established pipeline for extracellular vesicle small RNA data analysis [33]. Briefly, ExceRpt first automatically identifies and removes known 3’ adapter sequences. Afterwards the pipeline aligns against known spike-in sequences used for library construction, filters low-quality reads and aligns them to annotated sequences in the UniVec database and endogenous ribosomal RNAs. Reads that were not filtered out in the pre-processing steps are then aligned to the mouse genome and transcriptome using STAR aligner [34]. The annotations were performed in the following order: miRbase, tRNAscan, piRNA, GENCODE and circRNA. Reads mapped to miRNAs were combined from sequencing obtained before and after size-selection and were corrected for batch effect using RUVSeq [35]. Normalization factors were calculated using the TMM [36] method and differential expression was performed using the edgeR package [37] in R.

For cumulative distribution plots, miRNA targets (all and conserved) were downloaded from TargetScan release 7.2 [38]. When using the context++ scores, targets were split into 3 same-frequency groups according to their scores. *P*-values were calculated using a Kolmogorov-Smirnov test between the first and the last groups (i.e., strongest and weakest targets). The miRNA pathway analysis was conducted using a web-server tool DIANA-miRPath [23], where targets were predicted-derived from DIANA-TarBase v6.0, a database of experimentally validated miRNA targets. The adjusted *P* cutoff value of 0.05 was used for the identification of expressed pathways. The miRNAs and their corresponding target pathways information was extracted and plots were generated in R. ggplot2 [39] and ComplexHeatmap [40] R packages were used for generation of figures.

## Data availability

The datasets collected for this study are available as follows:

1. sRNA-seq dataset of epididymosomes before size-selection :
2. sRNA-seq dataset of epididymosomes after size-selection :
3. Codes used for bioinformatics analysis of the RNA-sequencing datasets:

## Acknowledgements

We thank Pierre-Luc Germain for advice on data analysis and for generating cumulative distribution plots, Irina Lazar-Contes for help with MSUS breeding, Silvia Schelbert for work on the animal license, Emilio Yandez at Function Genomics Center Zurich (FGCZ) for advice on the small RNA sequencing, Alekhya Mazumkhar for help with nanoparticle-tracking analysis, Yvonne Zipfel for animal care, Zurich Integrative Rodent Physiology facility for performing cholesterol measurements. We also thank Eloise Kremer, Ali Jawaid and Mea Holmes for their initial contributions to the project. The Mansuy lab is funded by the University Zürich, the Swiss Federal Institute of Technology, the Swiss National Science Foundation (31003A-135715), ETH grants (ETH-10 15-2 and ETH-17 13-2), the Slack-Gyr foundation, the Escher Foundation. Deepak K. Tanwar is supported by the Swiss Government Excellence Scholarship. Martin Roszkowski was funded by the ETH Zurich Fellowship (ETH-10 15-2).

## Authors contributions

AA and IMM conceived and designed the study. FM and MR performed the MSUS breeding and collected tissue samples. AA and DKT performed data analysis and generated figures. AA wrote the manuscript with input from DKT and IMM. AA performed all experiments for RNA sequencing and all molecular analyses. IMM supervised the project and raised funds.

## Supplementary Figure Panels

**Supplementary Figure 1.**
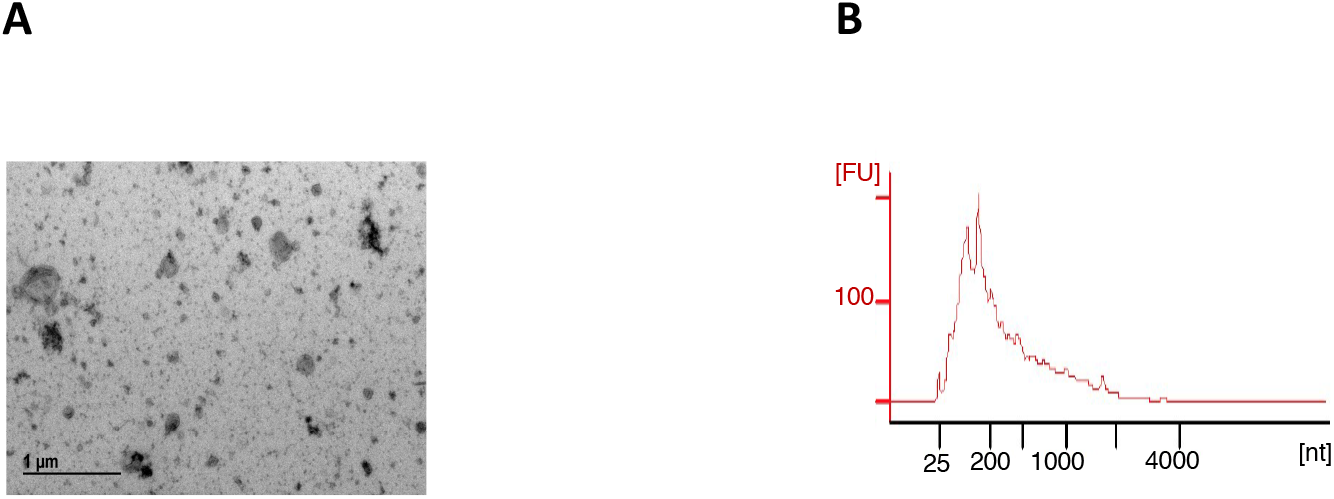
(**A**) Electron microscopy images of the preparations were used to access the size and heterogeneity of the isolated populations. (**B**) The RNA amount of epididymosomes preparations was assessed by high-resolution automated electrophoresis.

**Supplementary Figure 2.**
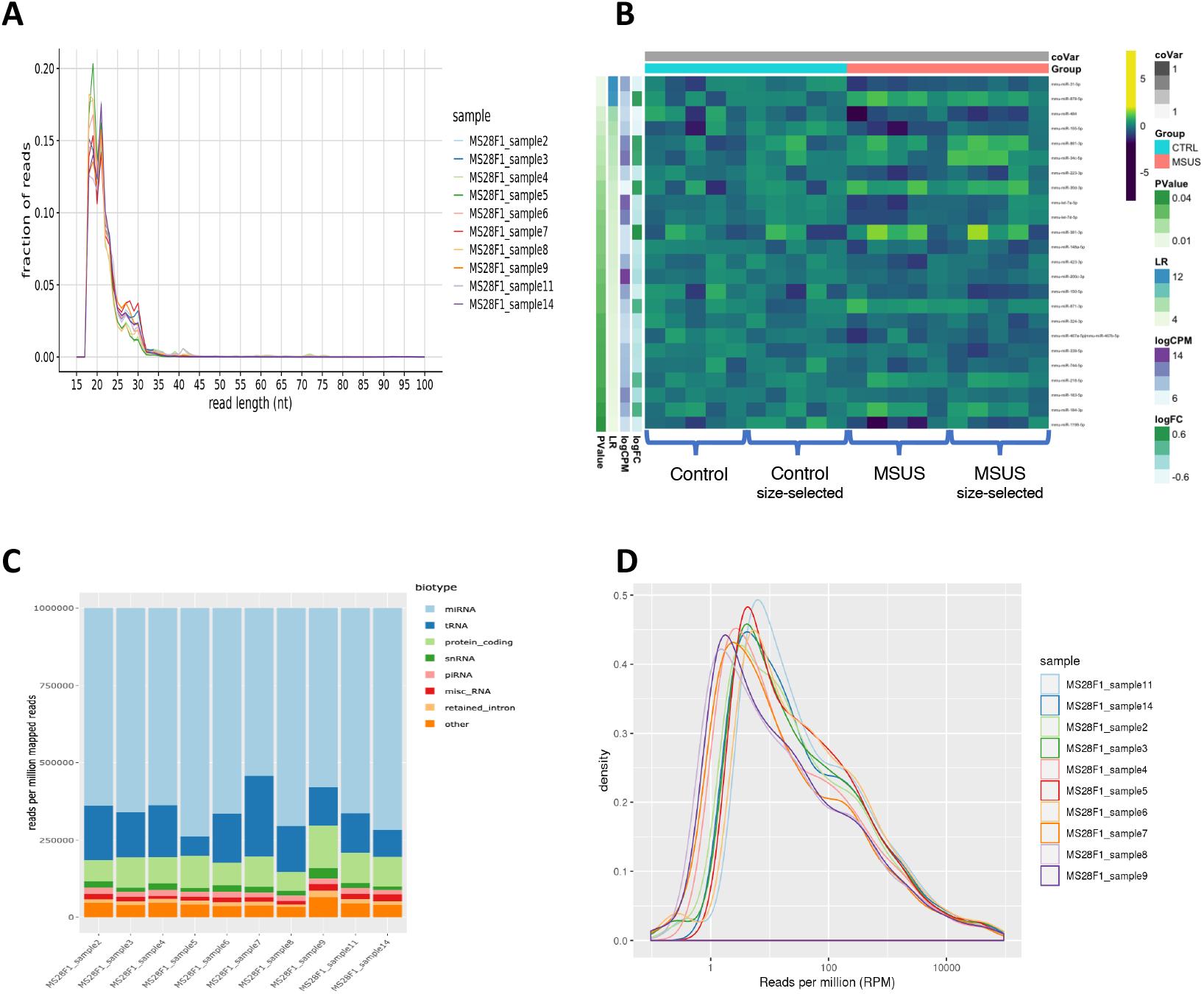
(**A**) RNA read-length distributions of cauda epididymosomal small RNAs after size-selection. (**B**) Heatmap of top miRNAs of cauda epididymosomal small RNA-seq before and after size-selection. Each row depicts a miRNA and each column depicts a sample. (N= 10 Controls, N=10 MSUS). (**C**) Representative distribution of RNA biotypes from cauda epididymosomal small RNA-seq after size-selection. (N = 5 Controls, N = 5 MSUS). (**D**) miRNA abundance distributions plot of cauda epididymosomal small RNA-seq. (N = 10 Controls, N = 10 MSUS)

**Supplementary Figure 3.**
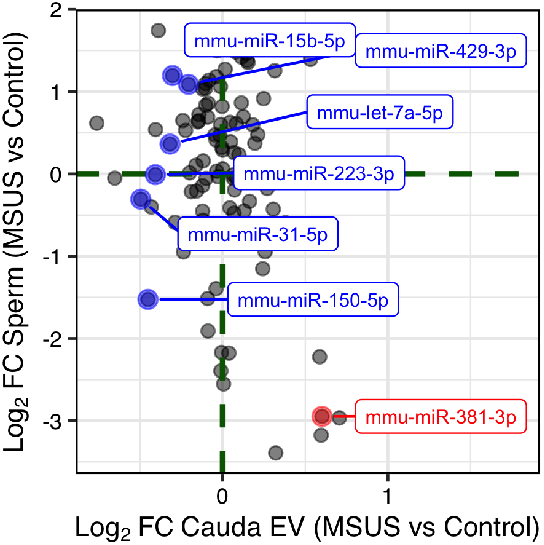
Scatterplot of log fold-change of miRNAs in sperm and cauda epididymosomes. miRNAs down-regulated (blue) and up-regulated (red) in MSUS cauda epididymosomes.

**Supplementary Figure 4.**
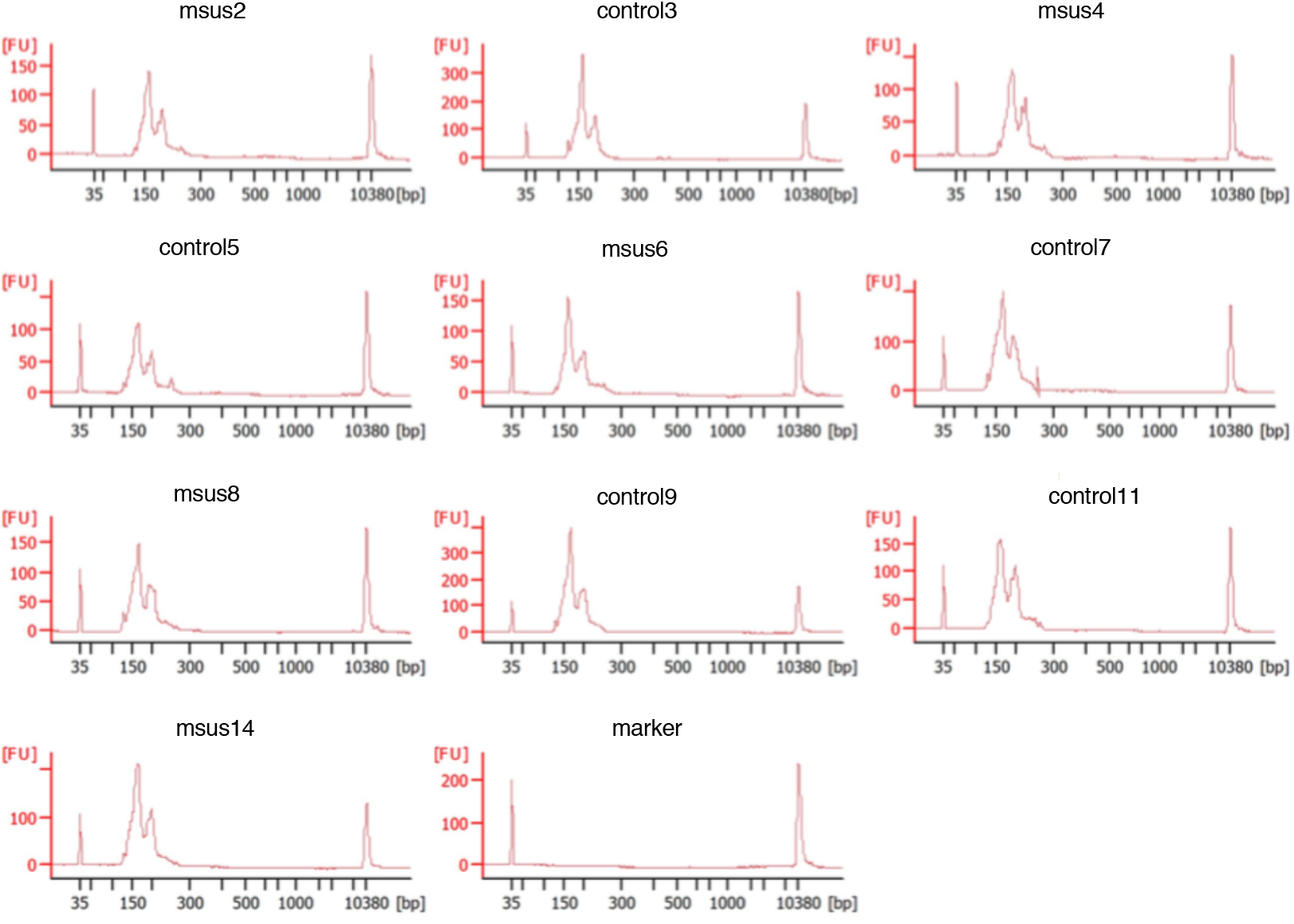
Profiles of cauda epididymosomal small RNAs by high-resolution automated electrophoresis. (N = 10 Controls, N = 10 MSUS).

**Supplementary Figure 5.**
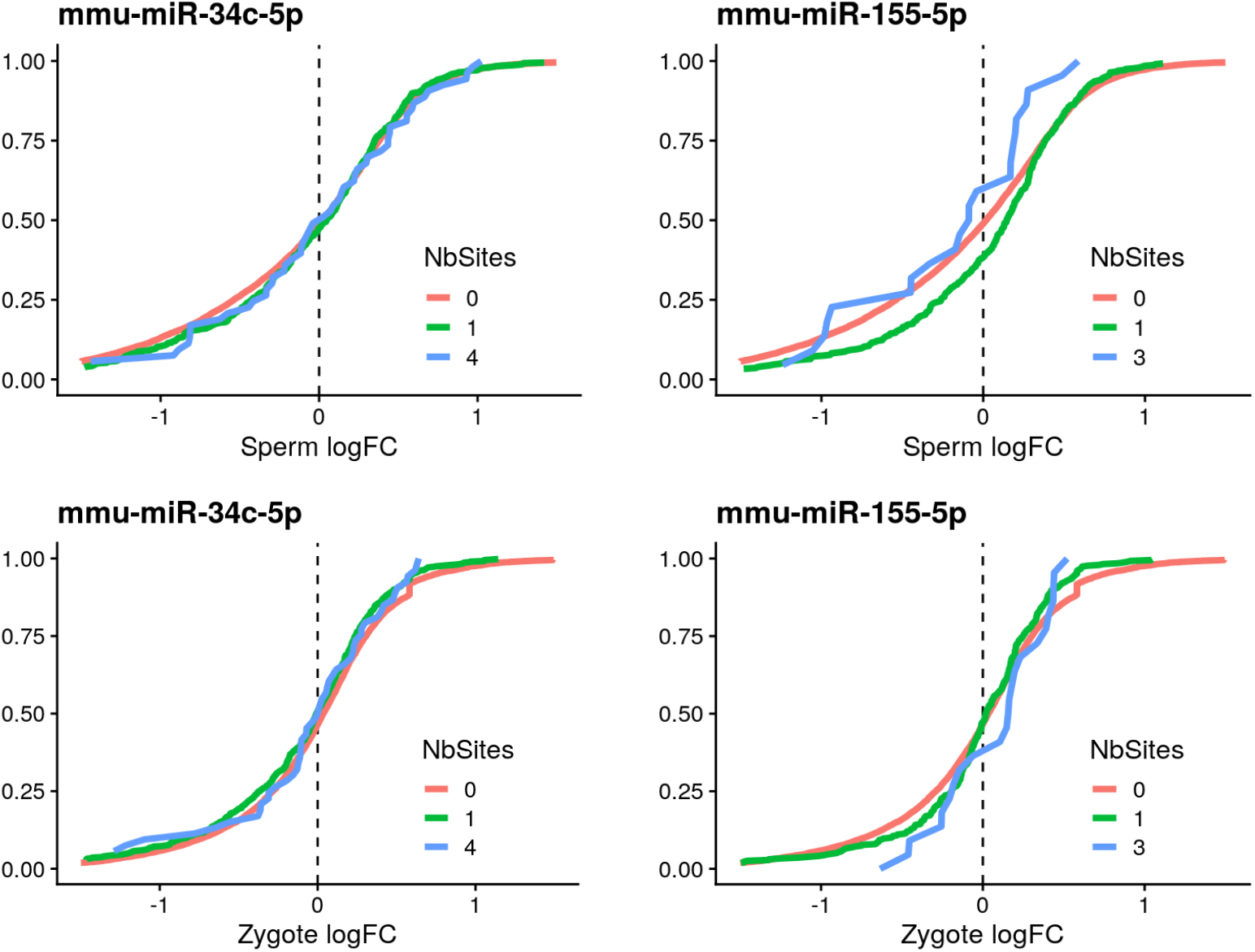
Cumulative distribution plots of miR-34c-5p, miR-155-5p targets in differentially expressed genes (P < 0.05) from MSUS sperm and zygote RNA-sequencing.

**Supplementary Figure 6.**
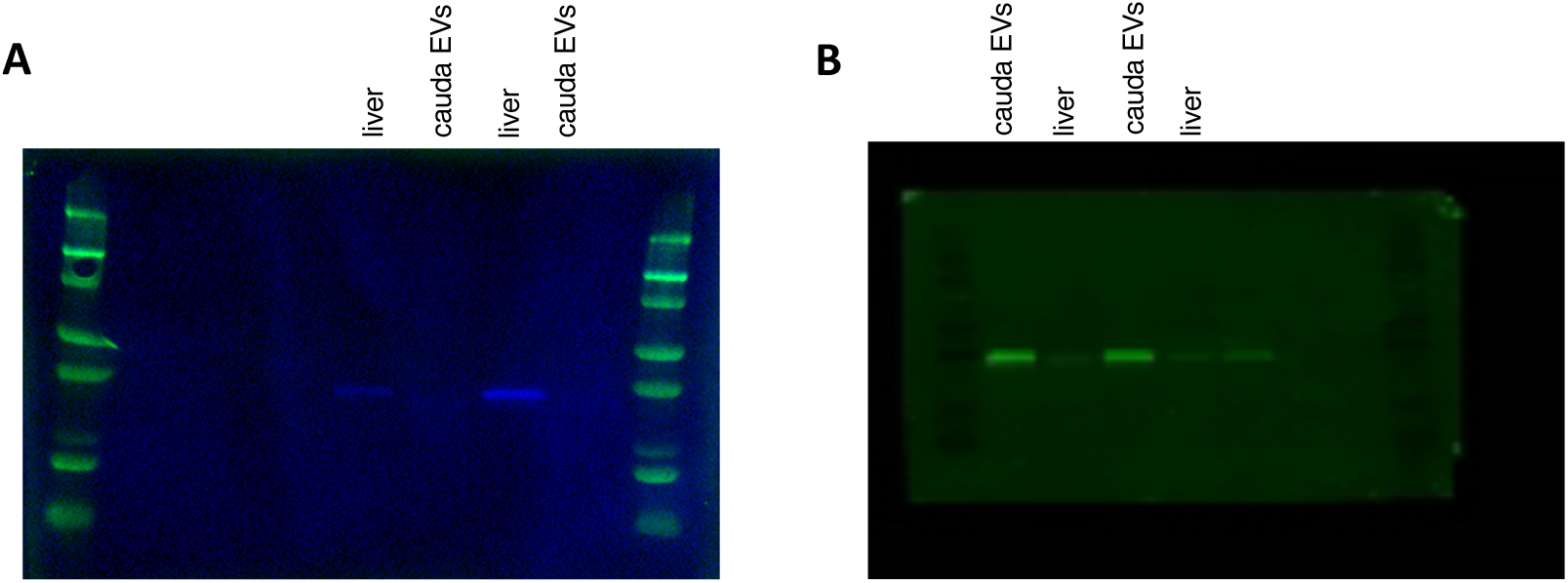
Immunoblot images of cauda epididymosomes and liver cell lysate, stained with cellular marker GAPDH (37kDa) and extracellular vesicle marker CD9 (25kDa). Same membrane was incubated and imaged separately.

